# Interpreting deep mutational scanning data resulting from selections on solid media

**DOI:** 10.1101/087072

**Authors:** Justin R. Klesmith, Sarah Thorwall, Timothy A. Whitehead

## Abstract

Deep mutational scanning is now used in directed evolution experiments to quantify the change in frequency of a cellular variant in a mixed population. A key concern is the extent to which the enrichment of a variant in a population corresponds to a fitness metric like relative growth rate or survival percentage. We present here analytical equations converting the enrichment of a variant to fitness metrics for plate-based selections. Using isogenic and mixed cultures we show that growth rates and survival percentages correlate for antibiotic plate-based selections. These results are important for proper interpretation of data resulting from deep sequencing.

## Introduction

Deep sequencing has emerged as a powerful, enabling tool for protein engineering (Jardine *et al.*, 2016, Klesmith *et al.*, 2015, Koenig *et al.*, 2015, Whitehead *et al.*, 2012). Deep sequencing-based measurements allow one to estimate the frequency of each mutational variant in a population to be screened or selected (Fowler *et al.*, 2014, Hietpas *et al.*, 2012). The end-point measurement is an enrichment ratio (ɛ_i_), defined as the base 2 logarithm of the frequency change of the variant i in the selected population compared to a reference population. A key question in such deep mutational scanning experiments is the extent to which this enrichment ratio corresponds to a fitness value or phenotype (Boucher *et al.*, 2016, Kowalsky *et al.*, 2015).

Many directed evolution experiments involve selections on solid media. Typically, a population of cells expressing the mutated protein of interest is plated on a solid support containing selective media, and the selected hits are colonies that survive or are larger than other colonies after a set amount of time. A major problem with plate-based selections is that the output is binary – a hit or not a hit. To remedy this, several groups have used deep sequencing to determine an analog fitness for each variant (Elazar *et al.*, 2016, Firnberg *et al.*, 2014, Hsiau *et al.*, 2015, Kim *et al.*, 2013) where the resulting fitness metric is usually normalized to the enrichment ratio of the starting sequence. However, the fitness metric calculated from enrichment ratios should depend strongly on whether the relative growth rates between variants differ, whether a variant survives the initial plating condition differentially, and the initial and final biomass concentrations. To that end, our objective in this work is to present analytical equations converting enrichment ratios to unambiguous fitness metrics for plate-based selections, and to supply experimental validation using an existing genetic selection for an antibiotic-based selection.

## Theory

On solid media, growth of bacterial biomass follows exponential behavior during the growth phase (Fujikawa and Morozumi, 2005). Importantly, the specific growth rate during exponential phase and the final density is independent of the plating density (Fujikawa and Morozumi, 2005). Therefore, for our model we assume that the growth model for each variant i can be written as:

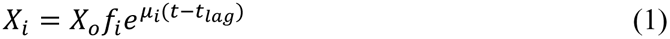

Where X_o_ is the initial biomass for variant i, X_i_ is the biomass of variant i evaluated at time t, t_lag_ is the lag time, µ_i_ is the specific growth rate, and f_i_ is the fraction of variant i that survives the initial selection at t=0. Based on work from van Heerden et al, we assume that the lag time is the same for all variants (van Heerden *et al.*, 2014). Equation (1) does not capture the characteristic sigmoidal shape of microbial growth curves – we neglect this for simplicity. We note that the inflection point on microbial growth curves typically occur within 1 or 2 average population doublings of the maximum biomass concentration. Thus, the error resulting from this simplification is likely to be minimal.

Using the above exponential growth model, it can be shown for conditions where t is greater than t_lag_ and X_i_ is less than X_max_ that the enrichment ratio of a given clone in the population can be determined by µ_i_ and f_i_:

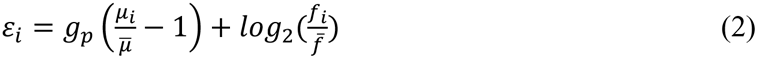

Here 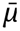and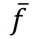 are the population-averaged values, and gp is the population-averaged number of doublings. Importantly, the above three parameters are experimentally measurable and, furthermore, common to all variants. Equation (2) shows that a given variant is enriched in the population if the specific growth rate is faster than the population average and/or if the fraction of surviving colonies is higher than the population average.

There are two limiting cases. If the fraction of growing cells is the same for each variant, as may be the case for selections coupling growth with flux through primary metabolism (Klesmith, Bacik, Michalczyk and Whitehead, 2015) then the fitness equation becomes (Kowalsky, Klesmith, Stapleton, Kelly, Reichkitzer and Whitehead, 2015).

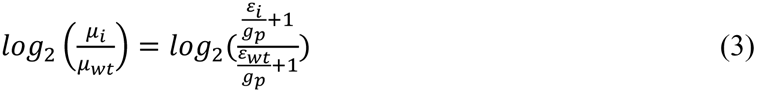

At the other limit, if the exponential growth rates are equivalent between all variants, the enrichment ratios can be normalized to a fitness metric defined as:

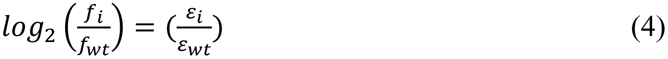

Here the wt subscript refers to the values from the wild-type sequence. While it is often implicitly assumed that equation (4) is the appropriate fitness metric for selections (Elazar, Weinstein, Biran, Fridman, Bibi and Fleishman, 2016), it could be the case that both the fraction of surviving variants and the relative growth rates vary between variants in the selection. In this case, time points must be taken over a time interval in order to calculate the specific growth rate and the surviving fraction for each variant.

To evaluate the appropriate form of the normalization expressions for antibiotic-based selections, we evaluated a genetic selection exploiting the twin-arginine translocation (TAT) pathway in Gram-negative bacteria (Fisher *et al.*, 2006). In this selection a protein of interest is fused between an N-terminal ssTorA Tat periplasmic export signal peptide and a truncated, active C-terminal TEM-1 beta-lactamase. Because the TAT pathway is thought to export only folded proteins, TEM-1 will be differentially exported to the periplasm based on the protein of interest in the fusion construct. Thus, mutations conferring enhanced fusion protein periplasmic localization can be selected on solid media containing different amounts of beta-lactam antibiotics.

## Results

We first evaluated selection-specific growth parameters for a ΔS4-A25 TEM-1 *bla* with the activity abrogating mutation S70A. We modified the TAT selection plasmid pSALECT-EcoBam (Addgene: #59705) (Hsiau, Sukovich, Elms, Prince, Strittmatter, Ruan, Curry, Anderson, Sampson and Anderson, 2015) by fusing a codon-swapped TEM-1 *bla* ΔS4-A25 in-frame after the XhoI restriction site. This active codon-swapped TEM-1 *bla* is used to avoid recombination with the N-terminal TEM-1 *bla* S70A. *E. coli* strain MC4100 harboring this plasmid was plated on LB-agar plates containing either 50, 100, or 200 µg/mL carbenicillin.

Colonies from a fresh transformation of *E. coli* MC4100 with the plasmid pSALECT-TEM-1(S70A)/csTEM-1 were used to start a culture in liquid LB media with 34 µg/mL chloramphenicol. This culture was grown at 30ºC at 250 rpm for 10 hours. Fresh LB agar plates with either 50, 100, or 200 µg/mL carbenicillin were made the day of the transformation and poured at a constant volume of agar per plate. The OD_600_ of the culture was measured after the 10 hour growth period to determine the cell density. The cells were diluted such that between 12-21 hours the cells were in exponential growth phase and plated. This initial plating density is different for each antibiotic concentration and was determined by control experiments. 150 µL of the diluted culture was spread onto the desired LB agar + carbenicillin plates. Plates were then placed into a humidified 30ºC incubator and grown for either 12, 15, 18, and 21 hours. At these time points a plate was taken out, and 1 mL of phosphate buffered saline was added onto the plate. All of the cells were scraped off of the agar plate and resuspended in this phosphate buffered saline. The final OD_600_ was then measured to determine the final cell mass. The specific growth rate was calculated from the natural log transformed cell densities at different time points. This experiment was repeated at least twice.

The fraction of surviving clones was determined by making serial dilutions of a culture with an OD_600_ of 1.0 ranging from 10^−3^ to 10^−7^ onto the selective LB agar + carbenicillin plates and LB agar + 34 µg/mL chloramphenicol plates. Plates were incubated at 30ºC in a humidified incubator and the number of distinct colonies recorded at each dilution. The fraction of surviving clones was calculated by dividing the number of distinct colonies on each carbenicillin plate by the number on the chloramphenicol plate. This was done with at least three biological replicates per antibiotic concentration.

For TEM-1 (S70A) the specific growth rate decreases at higher antibiotic concentrations (Figure 1a). This decreased growth rate correlates with low viability, although the growth rate plateaus at approximately half of the specific growth rate under conditions of high viability (Figure 1a). We performed the same experiment on a destabilized, catalytically inactive variant TEM-1 *bla* with mutations S70A, D179G (Wang *et al.*, 2002). As expected, the negative change in cellular viability is much greater than TEM-1(S70A) (Figure 1b). Similar to TEM-1(S70A), the growth rate decreases and plateaus to around half that of the high viability growth rate Figure 1b).

**Figure 1:**
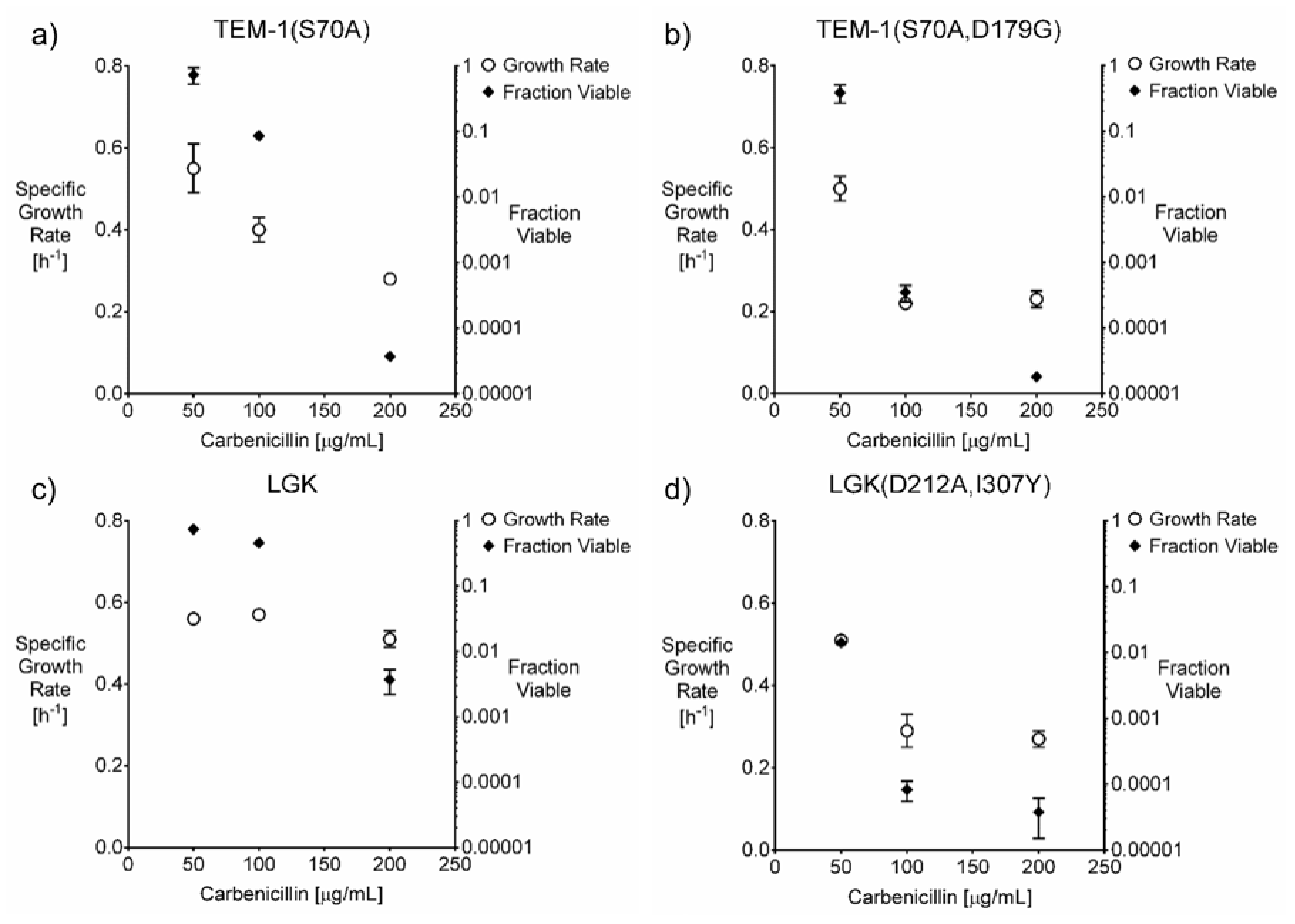
Specific growth rate (circles) and fraction viable (diamonds) of *E. coli* MC4100 expressing **a)** TEM-1(S70A), **b)** TEM-1(S70A,D179G), **c)** LGK, and **d**) LGK(D212A,I307Y). Error bars are 1 standard deviation of biological replicates (growth rates) or triplicates (fraction viable).

We performed the same set of experiments on a different enzyme system to confirm our initial observations. We used a codon optimized levoglucosan kinase (LGK) from *L. starkeyi* (Klesmith, Bacik, Michalczyk and Whitehead, 2015) (Figure 1c); and a destabilized, catalytically inactive variant LGK with mutations D212A, I307Y (Figure 1d). Similar results were seen for both strains where the growth rate decrease is independent of viability, the growth rate plateaus to half maximum for the destabilized variant, and viability has a more substantial decrease than growth rate. Therefore, both the growth rate and cellular viability impact enrichment ratios for plate based selections. From these results we predict that within a population the enrichment ratios for beneficial variants should increase with population size, and this effect should be accounted for in the analysis of deep mutational scanning experiments.

To evaluate gain of function mutations on a larger scale, we used nicking mutagenesis (Wrenbeck *et al.*, 2016) to create a single-site saturation mutagenesis library for residues 331-435 in LGK. We performed a TAT selection where we plated 100 µL of a culture of MC4100 *E. coli* at an OD_600_ of 1.0 with the LGK library on two 100 mm diameter petri plates with LB agar and 200 µg/mL carbenicillin per time point. Dilutions were also plated at 200 µg/mL carbenicillin and 34 µg/mL chloramphenicol to measure library viability. The plates were incubated in parallel at 30ºC in a humidified incubator, and the cellular mass was scraped and collected at time-points of 12, 14, 16, and 18 hours. The two replicates at each time point were pooled in equal volumes. The plasmids extracted from each time point were deep sequenced using an Illumina MiSeq in 300 bp paired-end mode using previously developed library preparation procedures (Kowalsky, Klesmith, Stapleton, Kelly, Reichkitzer and Whitehead, 2015). We then used Enrich (Fowler *et al.*, 2011) to calculate enrichment ratios for each variant at a given time-point relative to t = 0 hours. We observed 1,220 single point mutants with at least 15 read counts in the reference (unselected; t = 0 hours) population. Library statistics for the selections are shown in Table I. Processed deep sequencing datasets are freely available at figshare (www.figshare.com).

**Table I:** Library statistics, cellular densities, and fraction viable of time points.

|  |  |  |
| --- | --- | --- |
| Number of mutated codons | 105 |  |
| Reference sequencing reads post quality filter | 167,053 |  |
| Selected sequencing reads post quality filter |  |  |
| 12 hours | 110,898 |  |
| 14 hours | 184,418 |  |
| 16 hours | 186,130 |  |
| 18 hours | 182,375 |  |
| **Percent of mutant codons with:** |  |  |
| 1-bp substitution | 99.4 |  |
| 2-bp substitution | 84.2 |  |
| 3-bp substitution | 78.4 |  |
| All substitutions | 83.8 |  |
| **Percent of reads with:** |  |  |
| No nonsynonymous mutations | 35.0 |  |
| One nonsynonymous mutation | 52.4 |  |
| Multiple nonsynonymous mutations | 12.6 |  |
| **Coverage of possible single nonsynonymous mutations:** | 85.4 |  |
| **Biomass (OD<sub>600</sub>-mL):** |  |  |
| 0 hours | 0.1 | 0.1 |
| 12 hours | 26.2 | 28.4 |
| 14 hours | 43.1 | 42.8 |
| 16 hours | 52.0 | 58.0 |
| 18 hours | 75.4 | 85.2 |
| **Fraction viable:** |  |  |
| 12 hours | 0.0074 |  |
| 14 hours | 0.0126 |  |
| 16 hours | 0.0145 |  |
| 18 hours | 0.0153 |  |

For each variant we then plotted the enrichment ratio as a function of the observed average number of population doublings (Figure 2a). From this plot we extracted an enrichment ratio slope (slope_er_) as well as an average enrichment ratio (average_er_), defined here as the enrichment ratio at the midpoint of the best-fit linear regression line joining the four experimental data-points. The correlation coefficient between slope_er_ and average_er_ is only 0.38 (Figure 2b). However, there is a statistically significant relationship between the sign of slope_er_ and average_er_ (Fisher’s exact test binary classification p<0.0001). That is, if the slope_er_ is positive, the average_er_ is much more likely to be positive, and vice versa.

**Figure 2:**
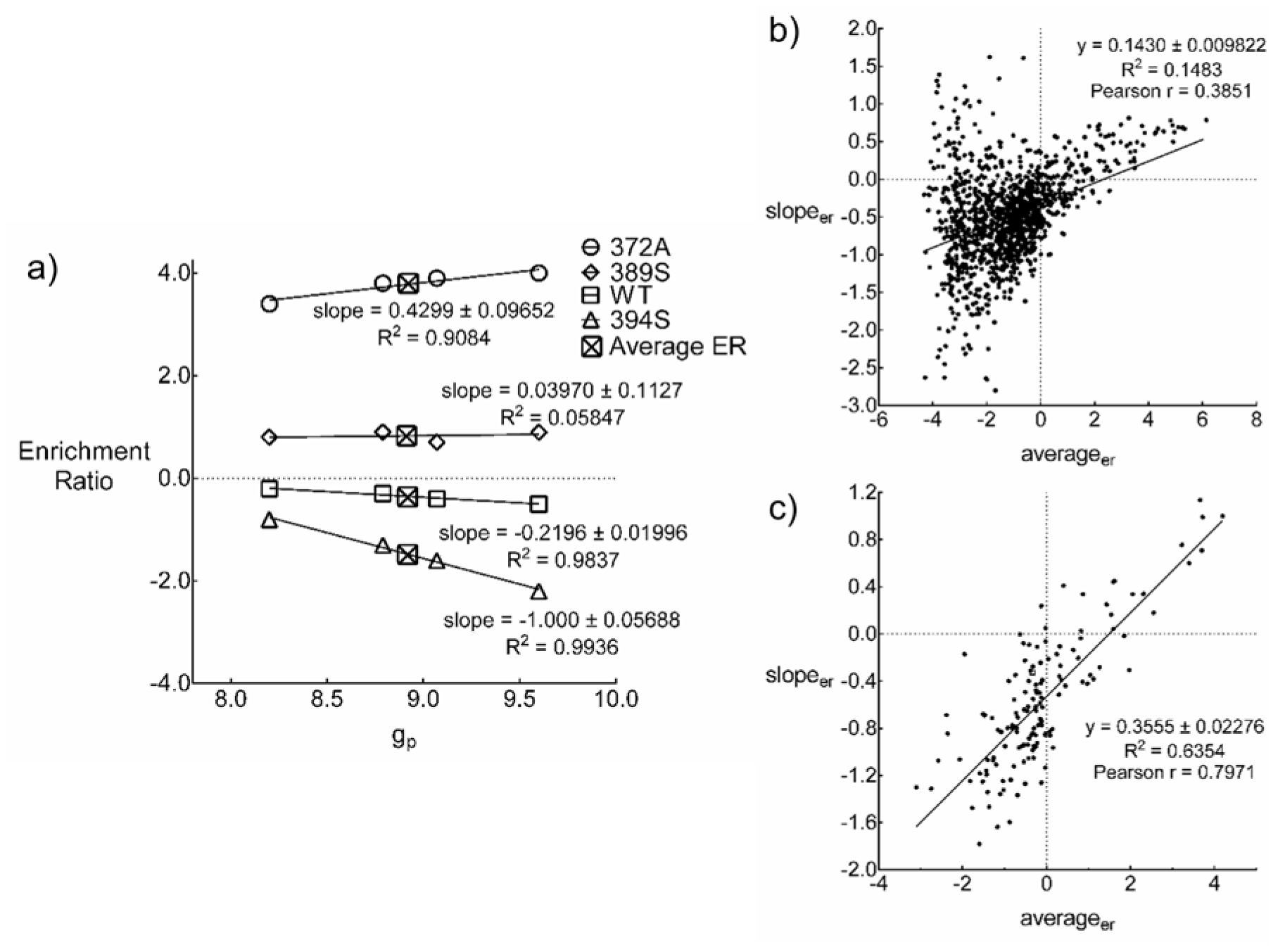
**a)** Enrichment ratio versus average population doublings of example variants showing an increase, neutral, or decrease in their enrichment over time. **b,c**: relationship between the change in enrichment ratio (slope_er_) and average enrichment ratio (average_er_) for **b**) all variants above 15 read counts; and **c**) all the subset of “high confidence” variants. Wild-type is indicated with an open square.

We reasoned that intrinsic counting error resulting from counting small numbers of variants could impact accurate determination of slopes. To test this assumption, we reran the above analysis on the subset of variants with over 100 read counts in the reference population (t = 0 hours) and at least 50 read counts on average in the four subsequent timepoints. For these resulting 142 variants, a much stronger relationship emerged between slope_er_ and average_er_ (Figure 2c), with the correlation coefficient now 0.80.

## Discussion

The above results show that enrichment ratios vary with respect to average number of population doublings for the plate-based TAT genetic selection in *E. coli*. We speculate enrichment ratios will vary for most coupled selections involving beta-lactam antibiotic resistance. For this particular antibiotic-based selection, these results are consistent with growth rate being a non-linear function of cell viability as shown here for isogenic cultures. While viability and growth rate may or may not be coupled for other types of plate-based selections, the above results have strong implications in the interpretation of deep mutational scanning data resulting from selections on solid media. In particular, implicit assumptions about the conversion of enrichment ratios to a fitness metric should be experimentally demonstrated. Furthermore, accurate determination of the slope is only apparent with variants well sampled in the population. As a practical matter, more depth of coverage for many deep mutational scanning experiments may be warranted.

## Acknowledgements

The authors would like to acknowledge L. Bergeron and members of the Whitehead lab in reading an earlier version of this communication. The authors declare no conflict of interest.

